# Aggregated and hyperstable damage-associated molecular patterns are released during ER stress to modulate immune function

**DOI:** 10.1101/652990

**Authors:** Alexander Andersohn, M. Iveth Garcia, Ying Fan, Max C. Thompson, Askar M. Akimzhanov, Abdikarim Abdullahi, Marc G. Jeschke, Darren Boehning

## Abstract

Chronic ER stress occurs when protein misfolding in the ER lumen remains unresolved despite activation of the unfolded protein response. We have shown that traumatic injury such as a severe burn leads to chronic ER stress *in vivo* leading to systemic inflammation which can last for more than a year. The mechanisms linking chronic ER stress to systemic inflammatory responses is not clear. Here we show that induction of chronic ER stress leads to the release of known and novel damage-associated molecular patterns (DAMPs). The secreted DAMPs are aggregated and markedly protease resistant. ER stress-derived DAMPs activate dendritic cells which are then capable of polarizing naïve T cells. Our findings indicate that induction of chronic ER stress may lead to the release of hyperstable DAMPs into the circulation resulting in persistent systemic inflammation and adverse outcomes.

## Introduction

The endoplasmic reticulum (ER) is the site of secretory and membrane-bound protein synthesis. Under conditions where ER protein synthesis rates exceed the folding capacity of the ER, unfolded or misfolded proteins accumulate in the ER lumen or membrane. The presence of an excess of misfolded proteins in the ER results in the activation of ER stress signaling pathways to restore homeostasis [1]. For example, ER chaperone content is increased while global protein synthesis rates are decreased in an effort to resolve the folding stress. If ER luminal protein folding stress cannot be resolved, pro-apoptotic pathways are activated resulting in cell death [2, 3]. Chronic ER stress is characterized by activation of this pathway without significant cell death resulting in cellular and organ dysfunction over extended time periods [4]. Inflammatory stimuli can lead to chronic ER stress in multiple cells and tissues [5]. For example, activation of the acute phase response results in dramatic upregulation in the synthesis of secretory proteins such as C-reactive protein resulting in hepatic ER stress [6, 7]. We previously found that chronic ER stress is prominent post-burn injury and persists for an extended period after the initial insult [8–17]. How chronic ER stress mechanistically contributes to post-burn inflammation and metabolic dysfunction is still unclear.

NLR Family Pyrin Domain Containing 3 (NLRP3) plays a central role in regulating inflammatory signaling transmitted by damage-associated molecular pattern molecules (DAMPs) derived from stressed or damaged cells [18, 19]. Inflammatory DAMPs include intracellular proteins such as high mobility group box 1 (HMBG1) and non-protein DAMPs such as nucleic acids, both of which are released from dying/damaged cells. Inflammasome activation by DAMPs leads to caspase 1 activation, resulting in maturation and secretion of IL-1β and downstream inflammatory responses [20, 21]. DAMP molecules are known to significantly contribute to systemic inflammation and adverse outcomes in trauma [22, 23]. We have previously shown that NLRP3 activation is central to post-burn responses including the induction of ER stress and systemic inflammation [17, 24, 25]. Pro-inflammatory cytokines such as IL-6 and IL-1β can lead to ER stress highlighting a positive feedback loop promoting inflammatory signaling [26–28]. Thus, DAMPs may contribute to systemic inflammation and long-lasting metabolic dysfunction after burn injury.

ER stress is known to induce the release of DAMPs. For example, chemotherapeutics can induce ER stress leading to the release of DAMPs and “immunogenic cell death” of cancer cells [29, 30]. It has also been shown that ER stress can lead to the release of DAMPs within secreted extracellular vesicles [31]. Here we show that inducing chronic ER stress in hepatoma cells leads to the release of non-vesicular DAMPs that are aggregated and hyperstable as determined by protease sensitivity. DAMP release was most likely associated with amphisome-mediated secretion *versus* apoptotic/necrotic cell permeabilization. The released DAMPs potently stimulated dendritic cell activation and the production of inflammatory mediators. Our results link chronic ER stress with the long-lasting inflammation and hypermetabolism found in trauma patients with significant therapeutic implications.

## Results

### ER stress leads to the secretion of intracellular proteins into the extracellular space

We previously demonstrated that ER calcium store depletion is a central mediator of post-burn hepatic ER stress [10]. To model this *in vitro* we depleted ER calcium stores with the SERCA pump inhibitor thapsigargin (TG) in HepG2 hepatoma cells, a well-characterized polarized human hepatocyte cell line model [32, 33]. We hypothesized that ER stress may lead to the release of aggregated proteins and/or extracellular vesicles into the media which could function as DAMPs. HepG2 cells were treated with TG for 24 hours and the media was subjected to differential centrifugation as in Figure 1A. There was a notable increase in the size of the cell-free high speed pellet in cells subjected to ER stress (Figure 1B). When the 40,000 *xg* supernatant and pellet fractions were run on SDS-PAGE, a large number of additional proteins were apparent in the pellet fraction of ER-stressed cells (Figure 1C). Identification of the bands by mass spectroscopy analysis revealed established DAMPs, such as histones, among other proteins which have not yet been established as *bone fide* DAMPs.

**Figure 1.**
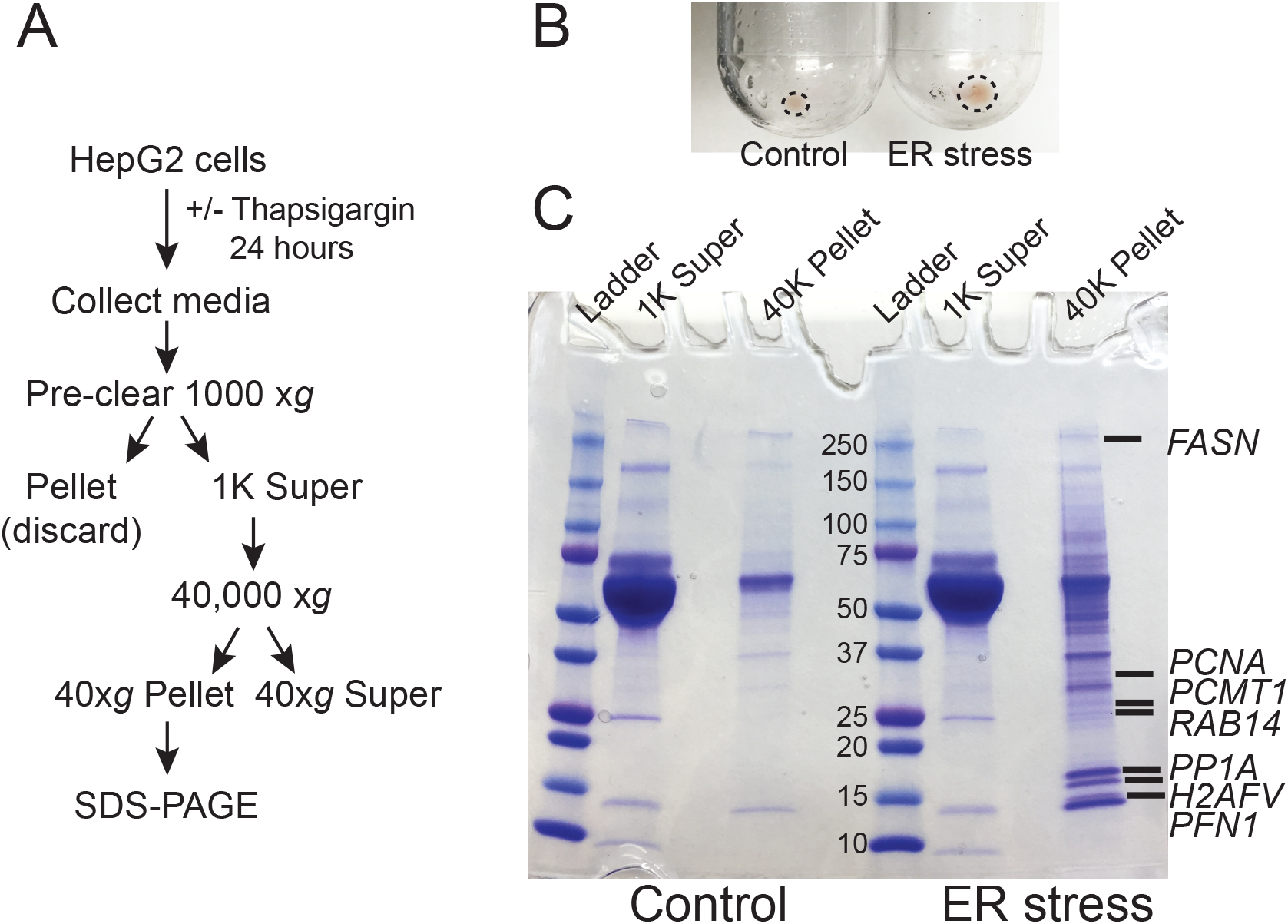
ER stress leads to the release of intracellular proteins. (A) Schematic of the treatment and purification protocol. (B) Image of the 40,000 *xg* pellets from a representative experiment. Pellet margins are highlighted. (C) Coomassie staining of the 1000 *xg* and 40,000 *xg* fractions. Fractions were run on a 4-20% SDS-PAGE gradient gel. The broad band running between the 50 and 70 kDa markers is bovine serum albumin from the media. Unique proteins identified by mass spectroscopy in ER stressed cells are indicated by gene name.

### Protein released during ER stress are non-vesicular and protease resistant

Recent evidence indicates that DAMPs may be secreted as extracellular vesicles during ER stress [31, 34]. To test whether the secreted proteins identified in Figure 1 are present within lipid vesicles, we subjected the 40,000 *xg* pellet to an additional purification step through a sucrose cushion. Using this protocol, vesicular components will remain in the sucrose cushion whereas high molecular weight non-vesicular (NV) components such as aggregated proteins will pellet through the cushion (Figure 2A; [35–37]). Using this fractionation protocol, there was a large prominent pellet present only in ER stressed cells (Figure 2B). When run on SDS-PAGE followed by Coomassie staining, the pellet fraction from ER stressed cells had an abundant number of proteins, some of which were also prominent in the 40,000 *xg* fraction (Figure 2C). Many of these proteins were also isolated in the NV fraction from another recent study which provided evidence that they are secreted in an amphisome-dependent manner [38]. Dilution of the sucrose cushion fraction and re-centrifugation to isolate extracellular vesicles resulted in no visible protein by Coomassie staining, indicating that extracellular vesicles are secreted at low levels in HepG2 under the conditions used in this study. Some of the proteins secreted from ER stressed cells are established DAMPs, such as histones, actin, and HMGB1. Western blotting confirmed the presence of these proteins in the NV fraction derived from the media of ER stressed cells (Figure 2D). We confirmed by Western blotting the presence of several other proteins which are not classically characterized as DAMPs such profilin-1 and enolase-1 (Figure 2D). ER stress leads to the production of misfolded and aggregated proteins which would be predicted to have increased resistance to protease digestion. To test whether proteins derived from the NV fractions were aggregated, we subjected these fractions to limited trypsin digestion. As a control we utilized total protein HepG2 Triton-X100 lysates. As shown in Figure 2E, almost all proteins present in HepG2 Triton-X100 lysates were digested within 15 minutes by *in vitro* trypsin digestion (Figure 2E). In sharp contrast, most proteins in the NV preps from ER stressed cells were detectable for the entire 180 minute time course (Figure 2F). Thus, ER stress leads to the secretion of known, and potentially novel, highly protease-resistant DAMPs which are not present within extracellular vesicles.

**Figure 2.**
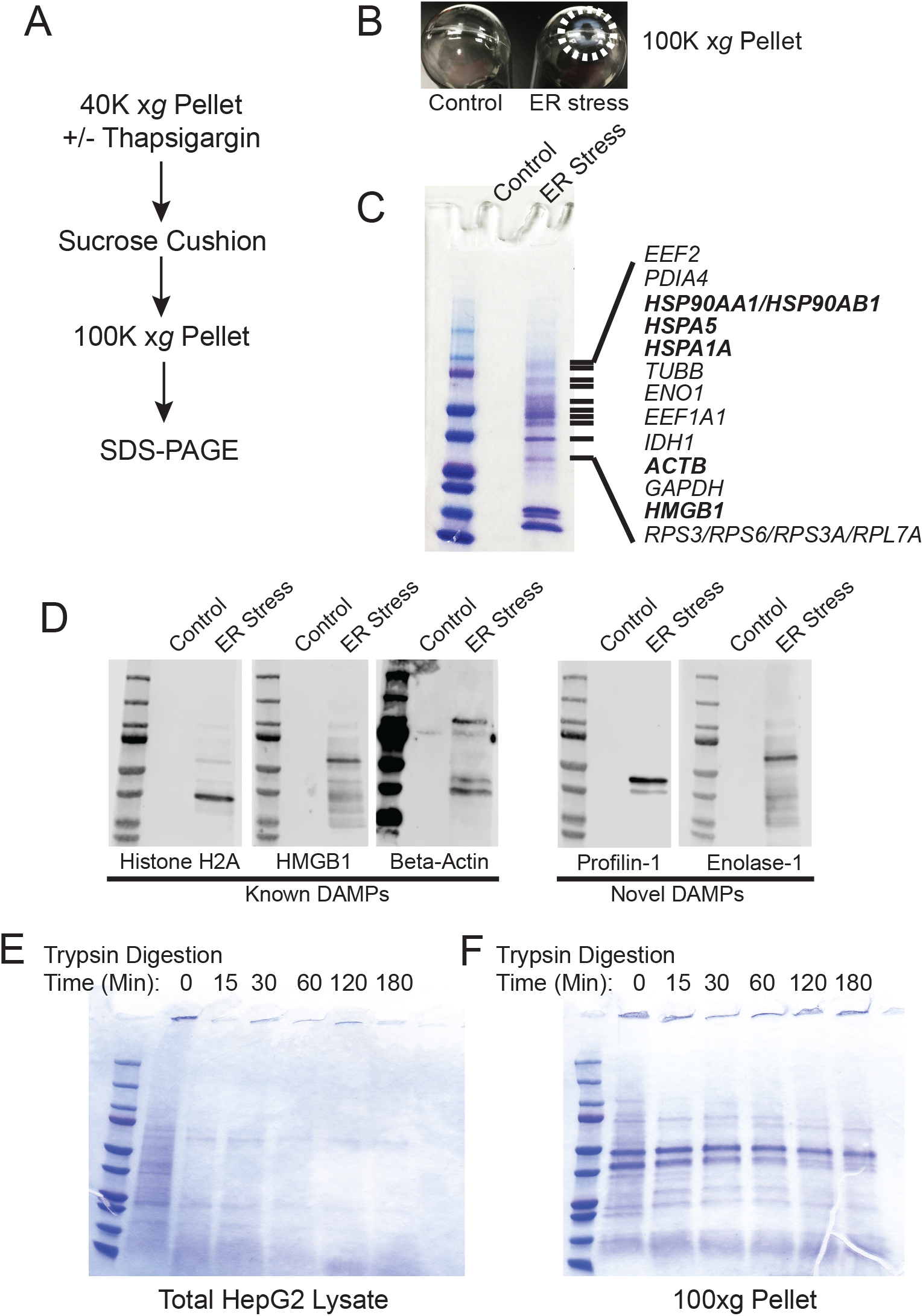
Proteins released from ER stressed cells are non-vesicular and protease resistant. (A) Schematic of the purification scheme. (B) Image of the 100,000 *xg* pellets from a representative experiment. Pellet margins are highlighted. Control (DMSO) treated cells do not have a visible pellet. (C) Unique proteins identified by mass spectroscopy in ER stressed cells are indicated by gene name. Known DAMPs are indicated in bold. (D) SDS-PAGE and Western blotting of a preparation as in (C) and identification with antibodies to the indicated proteins. (E) Trypsin digestion of HepG2 total cell lysate for the indicated times followed by Coomassie staining. (F) Trypsin digestion of the 100,000 *xg* pellet isolated from the media of ER stressed cells for the indicated times followed by Coomassie staining.

### Activation of apoptotic/necroptotic programs during ER stress

Release of NV proteins may either be through lysis of the plasma membrane or regulated release through other mechanisms such as the classical secretory pathway or non-canonical pathways such as the release of amphisome contents after fusion with the plasma membrane. To test these possibilities, we examined the activation of apoptotic and necrotic pathways in ER stressed HepG2 cells (Figure 3A). Thapsigargin dose-dependently induced the expression of BiP at all concentrations of TG after 24 hours of treatment, indicating activation of the ER stress program (Figure 3B). The broad spectrum kinase inhibitor staurosporine (STS), a positive control for the induction of apoptosis, did not induce BiP expression (Figure 3B). Light microscopy revealed significant cell loss only at TG concentration higher than 1 μM (Supplementary Figure 1). Consistent with this observation, significant cleavage of the caspase and calpain substrate α-fodrin was only obvious at TG concentrations of 1 μM and above (Figure 3C). To more quantitatively assess apoptosis induction, we measured enzymatic caspase-3 activities in lysates from HepG2 cells treated with either TG or STS. Concentrations of TG between 100 nM and 10 μM significantly activated caspase-3, however at much lower levels than the classic apoptosis inducer STS (Figure 3D). Propidium iodide (PI) is a cell-impermeant DNA dye commonly used to evaluate cell membrane permeabilization in apoptotic/necrotic models. Surprisingly, the number of PI positive cells in TG treated cells was higher than in STS treated cells at all concentrations except 10 nM (Figure 3E). One possible interpretation is that TG activated necroptotic signaling resulting in membrane permeabilization. However, there was no evidence of necroptosis activation as determined by phospho-MLKL Western blotting (Figure 3F-I). Single cell imaging of PI stained cells revealed TG treated cells did not have canonical nuclear staining, but rather display peripherally associated DNA staining (Supplementary Figure 2). We interpret this finding to indicate the amphisome-mediated secretion of free nucleic acids as seen by others [38] which are then stained by extracellular propidium iodide. Future work will be needed to confirm this interpretation. Regardless, it is possible to readily purify biochemically characterizable NV fractions from less than 100 mls of media of HepG2 cells treated with doses of TG as low as 100 nM. This concentration of TG leads to minimal activation of apoptotic/necroptotic signaling pathways, suggesting a secretion-based mechanism for NV protein release.

**Figure 3.**
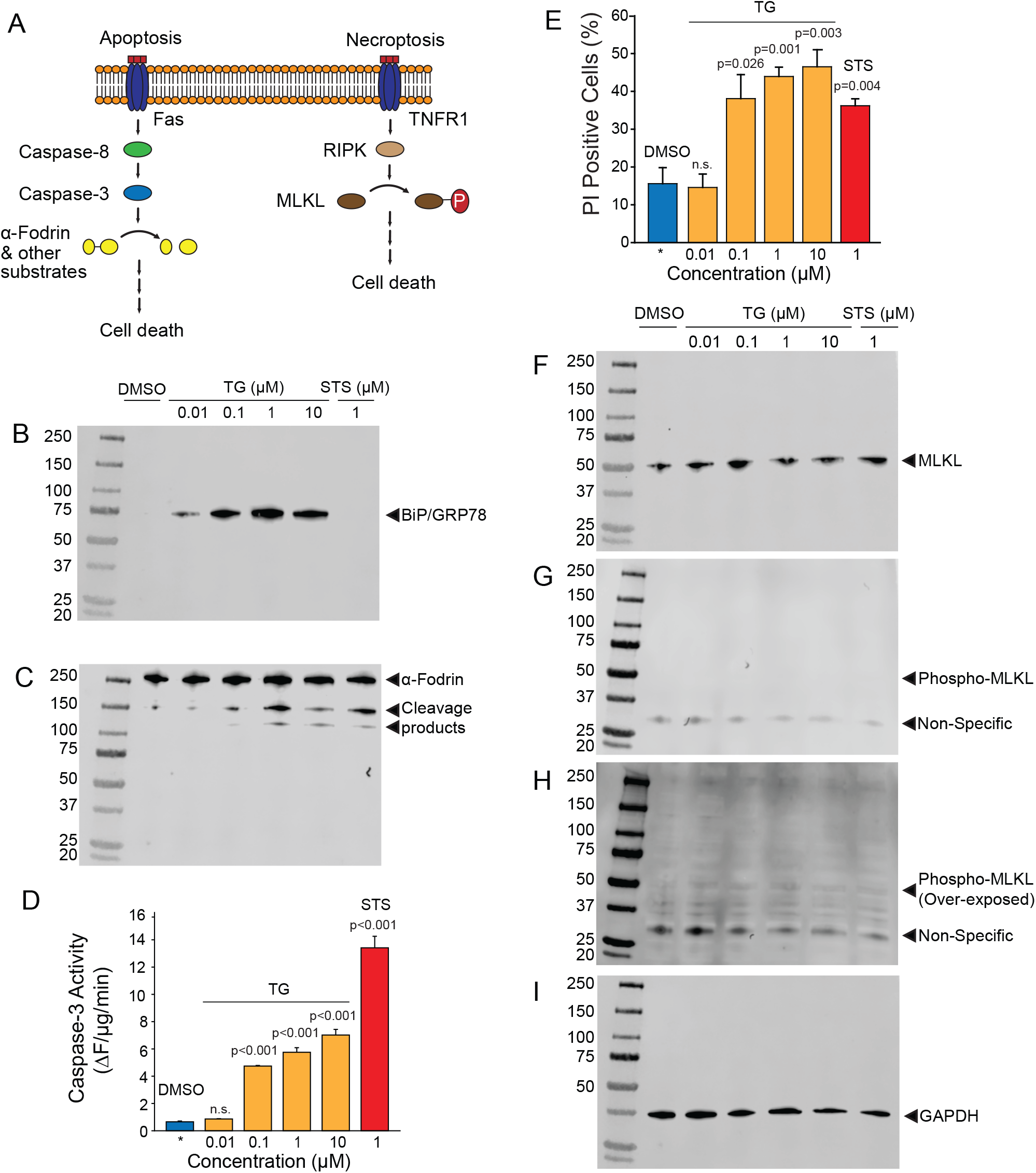
Activation of apoptotic and necroptotic pathways in TG treated HepG2 cells. (A) Schematic of apoptotic and necroptotic signaling pathways. (B) Western blotting of HepG2 lysates with BiP/GRP78, an ER stress marker, after the indicated treatments for 24 hours. TG, thapsigargin; STS, staurosporine. (C) Western blotting of the same membrane in (B) with α-fodrin, a marker of caspase and calpain activation. (D) Caspase-3 enzymatic activity in cells treated with the indicated compounds. Data is the mean +/- s.e.m. from three separate determinations. P values are indicated above the bars. (E) Propidium iodide (PI) positive cells treated with the indicated compounds. Data is the mean +/- s.e.m. from three separate blinded determinations. See text for additional information regarding scoring. (F-H) Western blotting of HepG2 lysates with MLKL and phospho-MLKL, a necroptosis marker, after the indicated treatments for 24 hours. (I) Same blot as in (F-H) probed with GAPDH as a loading control.

### ER stress-derived DAMPs regulate the expression of costimulatory molecules and cytokine production in dendritic cells

The NV fractions purified from ER stressed HepG2 cells contain well-characterized DAMPs (Figures 1–2). To test whether the NV fraction from ER stressed cells functions as a *bone fide* immune modulator, we tested whether this fraction could dose-dependently promote the maturation and activation of dendritic cells (DCs) [39]. During the development from bone marrow derived monocytes to DCs, there is a loss of macrophage marker F4/80 and increased expression of CD11b and CD11c (Figure 4A-B) [40]. To determine whether the putative NV-derived DAMPs shape DC phenotypes, we stimulated immature DCs with increasing concentrations of DAMPs purified from HepG2 cells treated with 100 nM TG. As a positive control we utilized 100 ng/mL lipopolysaccharide (LPS) and as a negative control we utilized PBS (vehicle). About 50% of DAMP-treated DCs were MHCII+, a marker suggesting they were ready to present antigens (data not shown). As shown in Figure 4B, DCs had increased expression of activation marker CD40 and CD86 as a function of DAMP concentration. Vehicle (PBS) treated DCs had no expression of CD40 or CD86. LPS is a well-known activator of DCs through TLR4. LPS treated DCs had strong expression of both CD40 and CD86. We next examined cytokine production in DAMP treated DCs. Both IL-6 and TNF-α production increased corresponding to DAMP concentration but in different manners. IL-6 production reached a plateau around 1 ng/mL when treated with 100 ng/mL DAMP. In contrast, TNF-α production kept rising as DAMP concentration was increased (Figure 4E-H). Although LPS induced stronger expression of CD40 and CD86, DAMP treatment induced similar levels of cytokine production when compared to LPS. These data demonstrate that the NV fraction purified from ER stressed HepG2 cells functions as a potent DAMP leading to the maturation of DCs and cytokine production.

**Figure 4.**
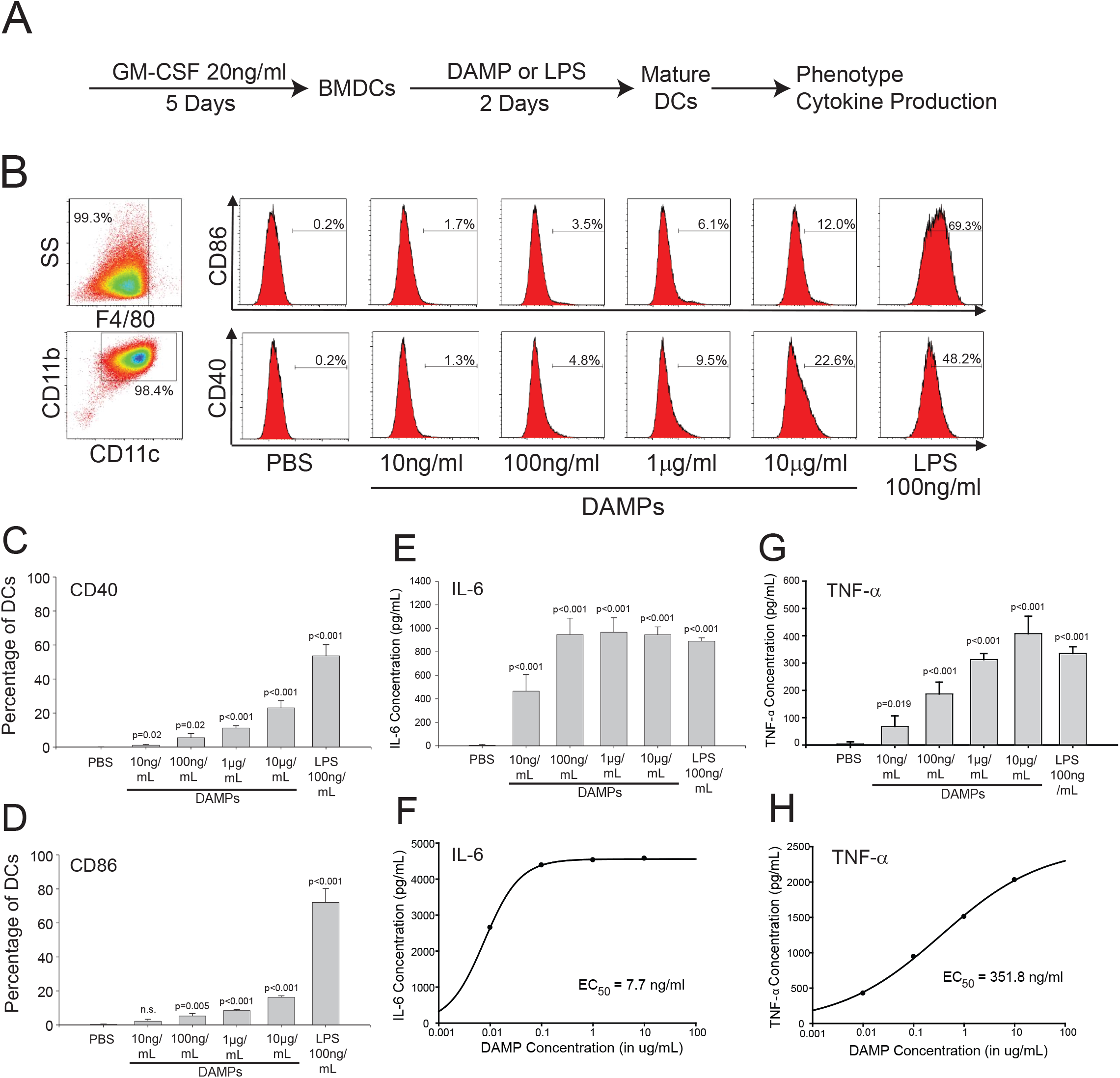
The non-vesicular (NV) fraction isolated from ER stressed cells activate dendritic cells as *bone fide* DAMPs. (A) Schematic of the *in vitro* bone marrow-derived dendritic cell (BMDC or simply DC) maturation protocol. (B) Macrophage and dendritic cell markers in control and DAMP treated BMDCs. Lipopolysaccharide (LPS) was used as a positive control. (C-D) Dendritic cell markers CD40 and CD86 after treatment with indicated compounds. Data are the mean +/- stdv from three separate determinations. (E-F) TNF-α cytokine production after treatment with the indicated compounds. (G-H) IL-6 production after treatment with the indicated compounds.

### Dendritic cells maturated by NV-derived DAMPs polarize naïve CD4+ T cells into a Th2 phenotype

Maturation of DCs through danger signals can be translated into the promotion of an inflammatory T-cell response. To further evaluate how NV-derived DAMPs modulate immune responses, DAMP-matured DCs were co-cultured with naïve CD4+ murine T cells for 5 days without additional DAMP stimulation. The co-culture supernatant was collected for cytokine analysis. After co-culture with DAMP-matured DCs, naïve CD4+ T cells produced a high amount of the Th2 cytokine IL-6 in a dose-dependent manner (Figure 5A). Furthermore, DCs treated with a relatively high concentration of DAMPs (10 µg/mL) induced T cells to produce another Th2 cytokine, IL-13 (Figure 5B). We also tested cytokines of other T helper cell phenotypes, however, no INF-γ or IL-17 production was detectable suggesting that the T cells were only differentiated into the Th2 phenotype. Thus, DCs were competent to present NV-derived DAMPs as a pro-inflammatory signal to T cells and could successfully induce a Th2 reaction. This has significant implications for systemic inflammatory responses in diseases associated with ER stress such as diabetes, cardiovascular disease, and trauma.

**Figure 5.**
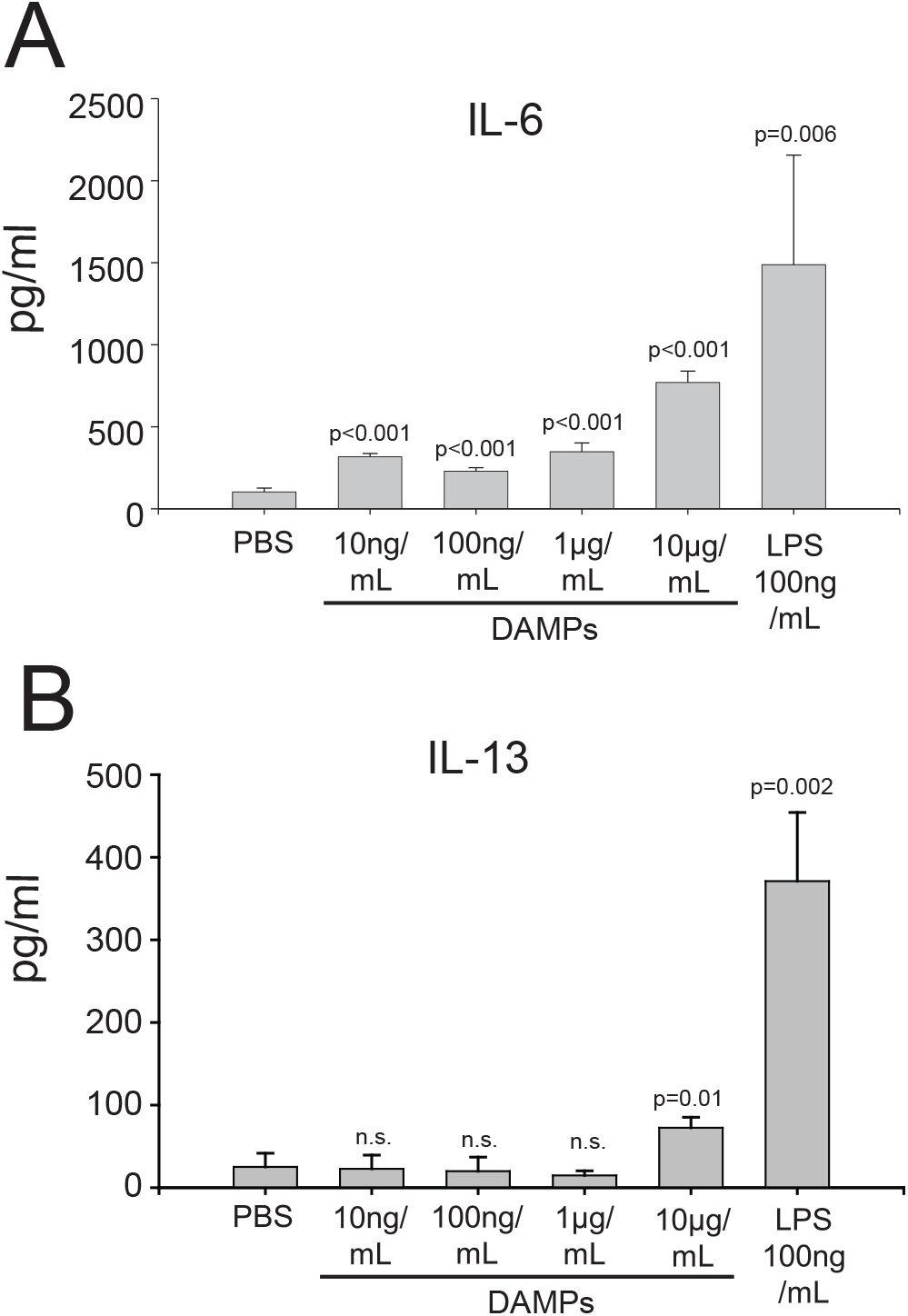
DAMP-differentiated dendritic cells are competent to differentiate naïve T cells into a Th2 phenotype. (A) Production of IL-6 in CD4+ T cells co-cultured with DAMP-stimulated DCs. The data represent the mean +/- stdv from three determinations. (B) Production of IL-13 in CD4+ T cells co-cultured with DAMP-stimulated DCs. The data represent the mean +/- stdv from three determinations. See methods for details.

## Discussion

In this report we show that inducing low levels of ER stress in HepG2 cells leads to the release of DAMPs in the context of minimal cell death. In our experimental model, the DAMPs released are not encapsulated with lipids in contrast to other models [31]. In a recent study on the secretion of intracellular components, it was found that many intracellular proteins and enzymes are secreted in an amphisome-dependent manner including cytosolic enzymes and nucleic acids [38]. The non-vesicular amphisome-secreted fraction isolated in this study has a protein composition remarkably similar to that observed in our study (Figures 1–2). Thus, we conclude that a similar mechanism mediates the release of intracellular components as DAMPs during ER stress. Future work will examine mechanistically whether TG-induced release of DAMPs requires amphisome formation and fusion with the plasma membrane.

Many studies have demonstrated the link between ER stress and the production of DAMPs. Cellular stress induced by chemotherapeutics cause the release of DAMPs and so-called “immunogenic cell death” or ICD [29, 30]. How ER stress couples to the release of DAMPs is still unknown, however it is thought to require cell membrane permeabilization. We found robust production of DAMPs which could be purified in biochemically characterizable amounts in cells stressed with low doses of TG. Under these conditions we found minimal caspase activation and cell permeabilization. Thus, we conclude there is active release/secretion of DAMPs during chronic ER stress. It is well established that ER stress activates autophagic pathways to rid the cell of excess unfolded/misfolded polypeptide chains [41–43]. We propose a similar model wherein chronic ER stress leads to activation of autophagic pathways to rid the cell of misfolded proteins. Autophagic vesicles then form amphisomes which fuse with the plasma membrane to release the excess of misfolded proteins in an effort to restore proteostasis. This is consistent with other recent studies showing the release/secretion of misfolded proteins in chronically stressed cells *in vitro* and *in vivo* [44, 45]. The released proteins then activate immune responses as DAMPs.

We have previously shown that burn injury leads to chronic systemic ER stress in multiple tissues, and in particular the liver [8–11, 13, 14, 17]. This contributes to metabolic syndrome leading to adverse outcomes. Mechanistically, burn injury leads to calcium store depletion *via* IP_3_R calcium channels leading to ER calcium store depletion and chronic hepatic ER stress [10]. The IP_3_R channel is a well-known central regulator of both ER stress and autophagic pathways [12, 46]. Thus, the IP_3_R calcium channel may provide the molecular link between burn injury, ER stress, and amphisome-mediated release of DAMPs. Future work will examine whether targeting this calcium channel is a potential therapeutic target to limit systemic inflammation and metabolic syndrome after a severe burn injury.

## Materials and Methods

### Animals

This study was carried out in accordance with the recommendations of the National Institutes of Health Guidelines for the Use of Laboratory Animals. The protocol was approved by the Animal Care and Use Committee of the University of Texas Health Science Center. Female C57BL/6 mice were bred at the animal facility at the University of Texas and used at the age of 8–12 weeks.

### Cell culture and treatment

HepG2 cells were purchased from ATCC and maintained in DMEM supplemented with 10% FBS, 1% penicillin/streptomycin, and 1% L-Glutamine. The cells were incubated at a constant temperature of 37°C and 5% CO_2_. When the cells had grown to 80%-90% confluency, thapsigargin (TG) purchased from Sigma Aldrich was added to fresh media at a final concentration indicated in the text/figures. Thapsigargin added at 100 nM and 1 µM for 24 hours produced similar amounts of NV DAMPs. Only DAMPs produced with 100 nM TG were used for biochemical characterization (Figure D-F) and DAMP functional analysis (Figures 4–5). An equal volume of DMSO was used as a control condition. A typical DAMP preparation required starting material from 60 mls media isolated from 2 x150 mm plates for each condition. This led to an average yield of approximately 50 µg protein at a concentration of 0.5 mg/ml.

### DAMP Isolation

The media collected from TG and DMSO treated cells were spun down at 1000 *xg* to remove dead cells and debris. The supernatants from this spin were transferred to round-bottomed centrifuge tubes and spun down at 40,000 *xg* at 4°C for 2 hours using an SS-34 rotor in a super-centrifuge. The pellets from the 40,000 *xg* spin were resuspended in an equal volume of PBS and 10 µL of each condition was removed for analysis by SDS-PAGE. The remainder of the resuspended pellets was diluted into PBS for density-gradient centrifugation. A 4 ml sucrose cushion consisting of 1M sucrose and 0.2M Tris base, pH 7.4 was transferred to centrifuge tubes and the resuspended pellets were carefully layered on top. The samples were placed in a Beckman SW-28 rotor and centrifuged at 100,000 *xg* at 4°C for 75 minutes in an ultracentrifuge. The pellets from this final centrifugation were resuspended in TTB buffer (120 mM KCl, 50 mM Tris/HCl pH 8.0, 1 mM EDTA, 1 mM DTT, 1% Triton-X100) or PBS and passed through a 25⅝ gauge needle before 10 µL of each condition was removed for analysis by SDS-PAGE.

### Proteomic Analysis

Protein bands on Coomassie-stained SDS-PAGE gels were excised with a razor and subjected to mass spectrometry-based protein identification by the Clinical and Translational Proteomics Service Center, The Brown Foundation Institute of Molecular Medicine, The University of Texas Health Science Center at Houston. In some cases, bands identified by mass-spectroscopy were confirmed by Western blotting.

### Trypsin Digestion

Untreated HepG2 cells were cultured for 48 hours, rinsed with PBS, and lysed with TTB buffer. The whole cell lysate was then diluted to the concentration of the DAMP isolation so that equivalent levels of protein were used. The whole cell lysate and DAMP isolation were then exposed to a trypsin solution (10 µg/mL trypsin, 30 µg/mL chymostatin, 100 µg/mL tosyl phenylalanyl chlormethyl ketone) in TTB for 0 min, 15 min, 30 min, 60 min, 120 min, 180 min at 37°C. At the end of the exposure time, the samples were quenched and run on separate gradient gels for each condition and Coomassie stained.

### In Vitro Generation of DCs and DC-naïve CD4+ T cell co-culture

Dendritic cells (DCs) were generated *in vitro* as previously described [1]. Briefly, tibias and femurs of C57BL/6 mice were removed under sterile conditions. Both ends of the bone were cut off, and the needle of a 1-mL syringe was inserted into the bone cavity to rinse the bone marrow out of the cavity. The cells were resuspended with Tris-NH_4_Cl red blood cell lysis buffer to remove the red blood cells. Bone marrow cells were then cultured in DMEM with 10% FBS, glutamine, nonessential amino acids, sodium pyruvate, HEPES, and penicillin/streptomycin (complete medium) for 2 hours. Floating cells were discarded and adherent cells were kept in complete medium with granulocyte-macrophage colony-stimulating factor (GM-CSF) (20 ng/ml) for 5 days. Complete medium and GM-CSF were renewed every 2 days. Immature DCs were treated with different concentration of DAMPs for 2 days and the supernatant was collected for cytokines analysis. In some experiments, stimulated DCs were processed for flow cytometry. Alternatively, stimulated DCs were re-plated in 96-well flat-bottom plates alone (3 x 105 cells/0.2 ml well volume) or with autologous naïve CD4+ T cells at a ratio of 1:10 for 5 days. Naïve CD4+ T cells were collected from the spleen and lymph nodes of WT mice using naïve CD4+ T cell isolation kit (Miltenyi Biotech). Co-culture supernatant was collected for cytokine analysis at the end of day 5.

### Assessment of the cytokine profile

Concentrations of IFN-γ, IL-13, IL-6, IL-17 (R&D Systems) and TNF-α (Thermo Fisher) in DCs or DC/T-cell co-culture supernatants were measured by ELISA, according to the manufacturer’s recommendations, and expressed in picograms per milliliter. Results were expressed as mean ± standard deviation.

### Phenotype analysis

DC phenotype was evaluated by flow cytometry using a standardized protocol [2]. Cells were kept on ice during all the procedures. For the extracellular markers, cells were stained with CD11c-AF700, F4/80-PE-Cy5, CD11b-APC-eF780, MHCII (I-Ab)-APC, CD40-Ef450 and CD86-FITC (ThermoFisher). Detection of cell surface markers was conducted using a Beckman-Coulter Gallios Flow Cytometer (BD Biosciences, San Jose, CA, USA) and data were analyzed by Kaluza Analysis Software. Results were shown as mean ± standard deviation.

## Conflict of Interest

The authors declare that the research was conducted in the absence of any commercial or financial relationships that could be construed as a potential conflict of interest.

## Author Contributions

A.A, M.I.G, Y.F., M.C.T. and A.A. performed experiments. A.A., M.I.G., Y.F., M.C.T., A.A. and D.B. analyzed data and prepared figures. M.G.J. and D.B. conceived of the project. All authors contributed intellectually, wrote, and approved of the final manuscript.

## Funding

This work was supported by National Institute of Health grants R01GM081685 (DB), R01GM087285 (MGJ, DB), and R01GM115446 (AMA). The proteomic studies were supported in part by the Clinical and Translational Proteomics Service Center at The University of Texas Health Science Center at Houston.

---

**Supplementary figure 1.**
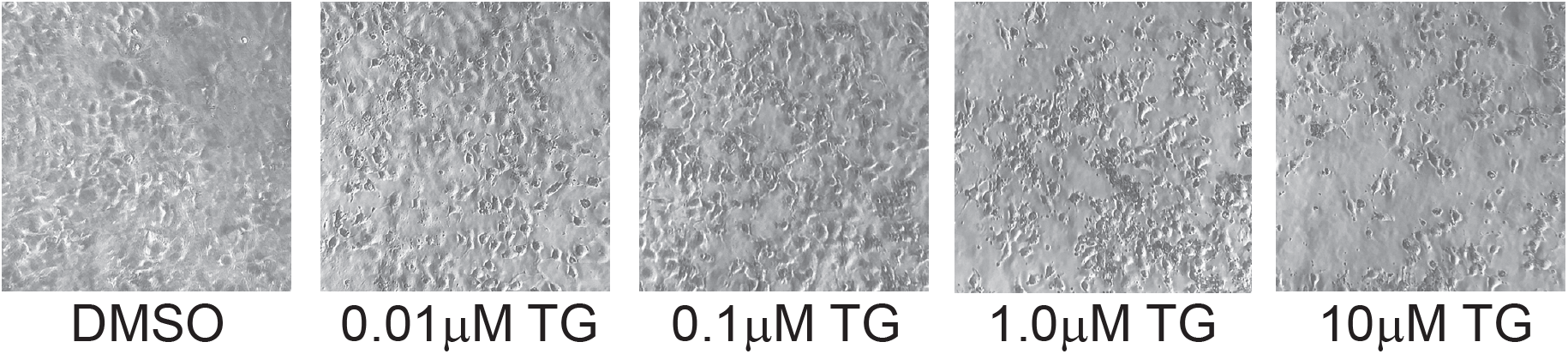
Phase contrast microscopy of TG treated HepG2 cells. Cells were treated with the indicated concentration of thapsigargin (TG) for 24 hours. Images are a representative field from at least three separate determinations.

**Supplementary figure 2.**
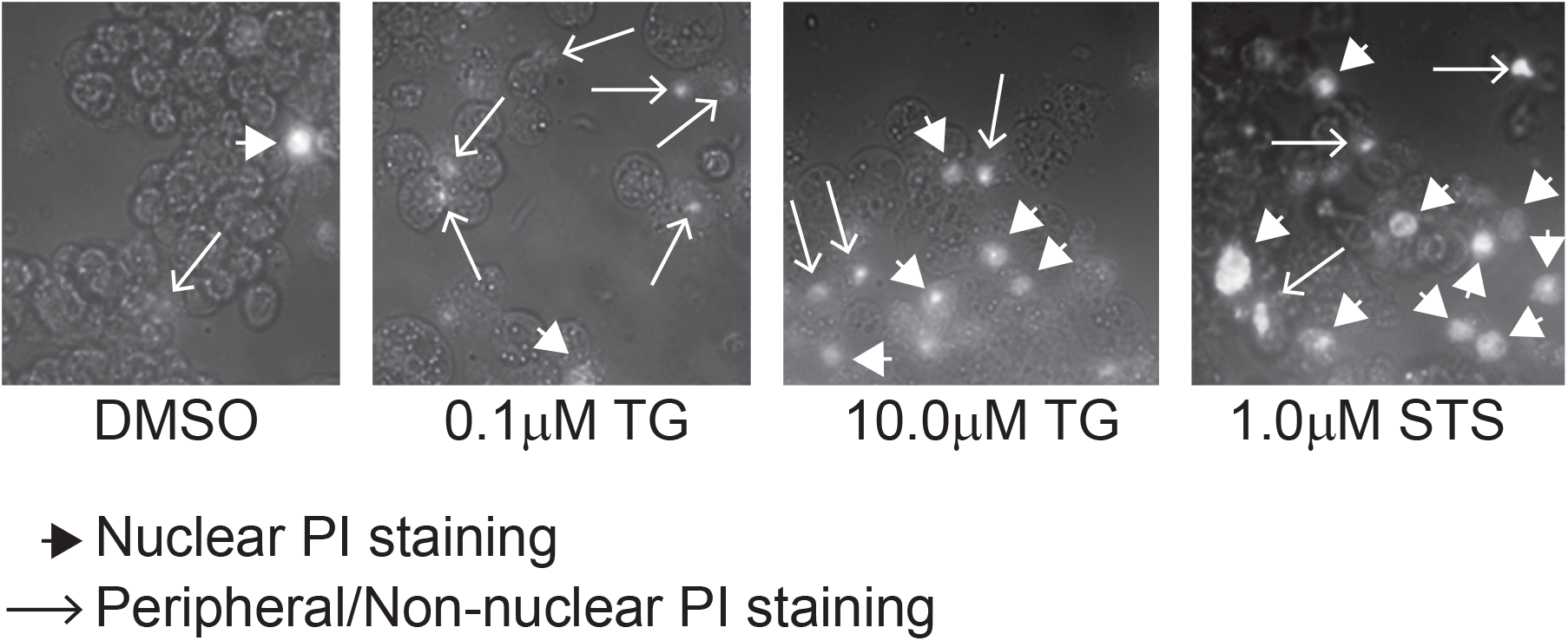
Propidium iodide (PI) staining of TG treated HepG2 cells. Cells were treated with the indicated concentration of thapsigargin (TG) for 24 hours. Images are a representative field from at least three separate determinations. Big arrow heads represent nuclear staining; narrow arrow indicate non-nuclear or peripheral DNA staining. Note staurosporine (STS) treated cells have abundant stained nuclei whereas ER stressed cells have more peripheral/non-nuclear staining.
See text for details.

